# How heterogeneity in glucokinase and gap junction coupling determines the islet electrical response

**DOI:** 10.1101/696096

**Authors:** J.M. Dwulet, N.W.F. Ludin, R.A. Piscopio, W.E. Schleicher, O. Moua, M.J. Westacott, R.K.P. Benninger

## Abstract

Understanding how cell sub-populations in a tissue impact the function of the overall system is often challenging. There is extensive heterogeneity among insulin-secreting β-cells within islets of Langerhans, including their insulin secretory response and gene expression profile; and this heterogeneity can be altered in diabetes. Several studies have identified variations in nutrient sensing between β-cells, including glucokinase (GK) levels, mitochondrial function or expression of genes important for glucose metabolism. Sub-populations of β-cells with defined electrical properties can disproportionately influence islet-wide free-calcium activity ([Ca^2+^]) and insulin secretion, via gap junction electrical coupling. However, it is poorly understood how sub-populations of β-cells with altered glucose metabolism may impact islet function. To address this, we utilized a multicellular computational model of the islet in which a population of cells deficient in GK activity and glucose metabolism was imposed on the islet, or where β-cells were heterogeneous in glucose metabolism and GK kinetics were altered. This included simulating Glucokinase gene (*GCK*) mutations that cause monogenic diabetes. We combined these approaches with experimental models in which *gck* was genetically deleted in a population of cells or GK was pharmacologically inhibited. In each case we modulated gap junction electrical coupling. Both the simulated islet and the experimental system required 30-50% of the cells to have near-normal glucose metabolism. Below this number, the islet lacked any glucose-stimulated [Ca^2+^] elevations. In the absence of electrical coupling the change in [Ca^2+^] was more gradual. As such, given heterogeneity in glucose metabolism, electrical coupling allows a large minority of cells with normal glucose metabolism to promote glucose-stimulated [Ca^2+^]. If insufficient numbers of cells are present, which we predict can be caused by a subset of *GCK* mutations that cause monogenic diabetes, electrical coupling exacerbates [Ca^2+^] suppression. This demonstrates precisely how heterogeneous β-cell populations interact to impact islet function.

**SIGNIFICANCE:** Biological tissues contain heterogeneous populations of cells. Insulin-secreting β-cells within the islets of Langerhans are critical for regulating blood glucose homeostasis. β-cells are heterogeneous but it is unclear how the islet response is impacted by different cell populations and their interactions. We use a multicellular computational model and experimental systems to predict and quantify how cellular populations defined by varied glucose metabolism interact via electrical communication to impact islet function. When glucose metabolism is heterogeneous, electrical coupling is critical to promote electrical activity. However, when cells deficient in glucose metabolism are in the majority, electrical activity is completely suppressed. Thus modulating electrical communication can promotes islet electrical activity, following dysfunction caused by gene mutations that impact glucose metabolism.

## INTRODUCTION

Most biological tissues are multi-cellular systems where a diverse range of cellular elements function collectively through dynamic interactions. Understanding the function of such a multicellular system requires understanding the characteristics of the constituent cell population and the way in which these populations interact over a defined architecture. As such the dynamics of multicellular systems often show complex emergent behaviors [1-4]. Many diseases can arise from genetic variations that impact molecular and cellular function. Given the complexities of multi-cellular systems, effectively predicting how molecular and cellular dysfunction lead to tissue and organ dysfunction, and cause disease is challenging.

The islets of Langerhans are micro-organs located within the pancreas that are critical for regulating blood glucose homeostasis. Complex multicellular dynamics occur between insulin-secreting β-cells and other endocrine cell types that underlie regulated hormone secretion. Death or dysfunction of β-cells that results in a reduction or absence of insulin secretion is the main cause of diabetes. Glucose metabolism and cellular excitability are key features of β-cell insulin secretion. Specifically, upon elevated blood glucose, glucose is transported into the β-cell and phosphorylated by the low affinity hexokinase, glucokinase (GK), which is the rate limiting step in glucose metabolism [5]. The increase in ATP/ADP inhibits ATP-sensitive K+ (K_ATP_) channels, causing membrane depolarization, activation of voltage-gated calcium channels and generating bursts of action potentials. The elevated oscillations in intracellular free-calcium activity ([Ca^2+^]) trigger pulses of insulin granule exocytosis [6-8]. In the absence of elevated ATP/ADP K_ATP_ channels remain open hyperpolarizing the cell, preventing voltage gated calcium channel activation and suppressing [Ca^2+^].

β-cells in the islet are electrically coupled by Connexin36 (Cx36) gap junction channels [9-11]. As a result of electrical coupling, β-cells exhibit coordinated [Ca^2+^] oscillations under elevated glucose and are uniformly silent under basal glucose [9,11-14]. β-cells are functionally heterogeneous, showing variations in electrical activity and insulin secretion [15-18], as well as varying gene expression profiles [19-21]. However as a result of electrical coupling, β-cells within the islet show near-identical [Ca^2+^] responses and oscillatory dynamics [11]. Nevertheless, the overall [Ca^2+^] response across the islet can be impacted by β-cell populations. In a normal islet nearly all β-cells are capable of responding to elevated glucose. However, a small number of inexcitable cells that remain hyper-polarized at elevated glucose can reverse this response and silence the whole islet [22-24]. Conversely a small number of highly excitable cells alone may be support the uniformly elevated [Ca^2+^] dynamics [25]. Understanding how different cell populations, characterized by different molecular signatures, interact and affect overall islet function is lacking.

We and others have previously examined how heterogeneity in K_ATP_ channel activity and electrical coupling impact overall islet [Ca^2+^] [9,22,24,26]. However, both single-cell based studies and in-situ analysis have demonstrated that β-cells exhibit substantial heterogeneity in glucose metabolism [27], which includes varying GK expression [21,25,28], metabolic fluxes [29-32], or mitochondrial function [19,21]. While extensive heterogeneity in glucose metabolism exists within the islet, it is not clear how cell sub-populations with differing GK expression or levels of glucose metabolism influence each other via electrical coupling.

Computational models have been utilized to examine how islet function is impacted by different cellular populations. For example, the frequency of islet [Ca^2+^] oscillations can be predicted by the relative numbers of slow-oscillating and fast-oscillating β-cells and their electrical coupling [33,34]. In the presence of a broad heterogeneous population, electrical coupling was also suggested to sharpen the islet response [34]. We predicted this concept computationally, where electrical coupling allows the islet to withstand a critical number of inexcitable cells and show near-normal glucose-stimulated [Ca^2+^]. Beyond this critical number, the islet sharply transitions to lack any glucose responsiveness [22,26], despite there being a population of β-cells that would otherwise be capable of responding to glucose. Computational models have also predicted that electrical coupling increases the impact of human K_ATP_ channel mutations that cause neonatal diabetes mellitus (NDM) [26]. These models predicted islet dysfunction was reversible upon modulating electrical coupling for many mutations, by allowing a population of β-cells to show [Ca^2+^] elevations. Mutations to *GCK* which reduce its activity can cause monogenic diabetes, either mature onset diabetes of the young (MODY) or NDM [35,36]. Prior computational studies therefore suggest that electrical coupling may play an important role upon heterogeneity to GK activity, including mediating how mutations to *GCK* impact islet function.

In this study we apply experimental and computational approaches to examine the role of gap junction mediated electrical coupling between β-cells in the presence of heterogeneity to GK activity. We examine how a defined population deficient in GK and glucose metabolism impacts islet function and how electrical coupling mediates the impact of this sub-population. We also examine how electrical coupling between a broad heterogeneous population of cells impacts islet function, and compare this with endogenous heterogeneity in the islet. Finally, we predict how human *GCK* mutations impact islet function, and the role that heterogeneity in glucose metabolism and electrical coupling play in mediating the impact of these mutations.

## MATERIALS AND METHODS

### Ethics Statement

All experiments were performed in compliance with the relevant laws and institutional guidelines and were approved by the University of Colorado Institutional Biosafety Committee (IBC) and Institutional Animal Care and Use Committee (IACUC, B-95817(05)1D).

### Animal Care

The generation of GK^lox/lox^ (Glucokinase with loxP sites flanking exon2); Pdx-Cre^ER^ (β-cell specific inducible Cre); and Cx36^−/−^ (Connexin36 global knockout) have been described previously [37-39]. Expression of variable GK deficiency was achieved by crossing GK^lox/lox^ and Pdx-Cre^ER^ mice, and inducing GK deletion in 8-24 week old mice by 1-5 daily doses of tamoxifen (50mg/kg bodyweight or 5mg/kg bodyweight as indicated) administered IP. Littermates lacking Pdx-Cre^ER^, lacking GK loxP sites, or wild-type C57BL/6 mice were used as controls. Mice were held in a temperature-controlled environment with a 12 h light/dark cycle and given food and water ad libitum.

### In-Vivo Measurements

Blood glucose was measured 3/week using a glucometer (Ascensia Contour, Bayer), and averaged over day 10-15 post-tamoxifen induction. Plasma insulin was measured at day 15 post-tamoxifen induction, from plasma extracted from blood samples centrifuged for 30 minutes at 3000 rpm at 4°C and assayed using mouse ultrasensitive insulin ELISA (Alpco).

### Islet Isolation

Pancreas dissection and islet isolation were performed in mice under Ketamine/Xylazine anesthesia that were subsequently euthanized via exsanguination and cervical dislocation or bilateral thoracotomy. Islets were isolated by collagenase injection into the pancreas through the pancreatic duct; the pancreas was harvested and digested, and islets were handpicked [40,41]. Islets were maintained in RPMI medium at 11mM glucose (Invitrogen) supplemented with 10% FBS, 100U/ml penicillin, 100μg/ml streptomycin, at 37°C under humidified 5% CO_2_ for 24-48 hours prior to study.

### Calcium Imaging

Isolated islets were loaded with 4 µM Fluo-4 AM (Invitrogen) for 45min at 37°C in imaging medium (125mM NaCl, 5.7mM KCl, 2.5mM CaCl_2_, 1.2mM MgCl_2_, 10mM Hepes, 2mM glucose, and 0.1% BSA, pH 7.4) and were imaged in 35mm glass bottom dishes maintained at 37°C. Fluo-4 fluorescence was imaged on a confocal microscope (Zeiss LSM800) with a 40× 1.2 NA water immersion objective, 488nm diode laser for excitation, and 490-550nm band-pass emission filter. Images were acquired at 1 frame/sec for 2min at 2mM glucose and for 5min at 11mM glucose after 15min of glucose stimulation. 20mM KCl was then added during continuous imaging. Microscope settings (integration time, scan time, gain, laser power) were constant for all images collected within the same day. For mannoheptulose (MH) experiments, after imaging islets at 11mM glucose, islets were imaged for 5min, after either 3, 5, or 10mM MH was applied for 15min at 11mM glucose.

### NAD(P)H imaging

NADH(P)H autofluorescence was imaged under two-photon excitation using a tunable mode-locked Ti:sapphire laser (Coherent Chameleon) set to 710nm for excitation. Fluorescence emission was detected at 425-475 nm using a non-descanned detector. A z-stack of 6 images were taken at 2µm intervals starting 10µm from the bottom of the islet. Images were acquired at both 2mM and 15min after 20mM glucose stimulation. Microscope settings (scan time, gain, laser power) were constant for all images collected within the same day.

### Image Analysis

All images were analyzed using custom MATLAB (MathWorks) scripts. For calcium imaging, a 5×5 smoothing filter was first applied. Areas with very low to no fluorophore staining were excluded from analysis by excluding pixels at or below the average fluorescence of a manually chosen background area. Saturated areas were also removed by limiting the area to intensities below the maximum. To determine if a cell was active, a peak detection algorithm was used to determine if areas of the islet had peak amplitudes greater than 1.5 times the average of quiescent reference cells [32]. A quiescent reference cell (or background region if no cells were quiescent) was chosen manually, and showed no significant fluctuations in intensity during the entire time course. The fraction of active area was then calculated as the number of pixels detected as active relative to the total number of pixels that were not excluded as background in the defined islet. Activity maps are presented in HSV format, with the hue representing the [Ca^2+^] oscillation synchronization, as described previously [11]. Briefly, cross-correlation coefficients (between 0-1) were calculated between every pixel area and a reference consisting of the average [Ca^2+^] fluorescence of the whole islet. This was calculated only for areas of the islet determined as active (see above). For MH treatments, [Ca^2+^] activity was normalized to the activity of the islet at 11mM glucose.

NAD(P)H images were analyzed by averaging the intensity across the z-stack for the islet. For each experimental condition these intensity values were normalized to mean NAD(P)H intensity at 2mM of islets from control mice that were acquired on the same day. The average intensity at 2mM was subtracted from the average intensity at 20mM to obtain the NAD(P)H response for the islet.

### Coupled β-cell Electrical Activity Model

The coupled β-cell model with and without stochastic noise was described previously [26] and adapted from the published Cha-Noma single cell model [42,43], with modifications to the ATP dependence as previously described [44]. All code was written in C++ and run on JANUS (University of Colorado Boulder) or Rosalind (University of Colorado Denver) supercomputers. Model code is included in the Supporting Information (Supplemental Information, File S1).

The membrane potential (*V*_*i*_) for each β-cell *i* is related to the sum of individual ion currents as described by [42]:

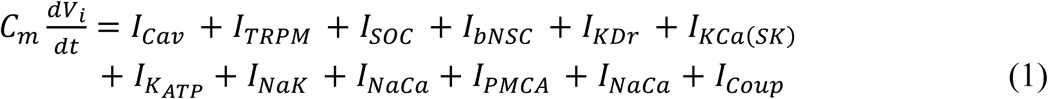

Where the gap junction mediated current *I*_*Coup*_ [22] is:

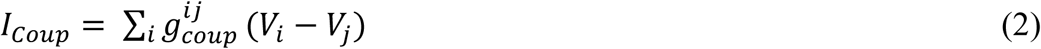

NADH concentration ([*Re*]) was calculated by solving the following differential equation [42]:

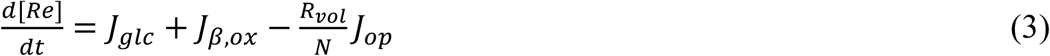

Where *J*_*glc*_ is the flux of glycolysis, *J*_*β, ox*_ is flux of β-oxidation and *J*_*op*_ is the flux of oxidative phosphorylation and ATP production. *N* =2.5 stochiometric conversion of NADH to ATP, and *R*_*vol*_ =2.5, the ratio of the volume of the cytosol to the mitochondria. ATP and ADP (bound (b) and free (f)) relationships have been previously described [42]:

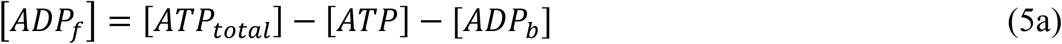

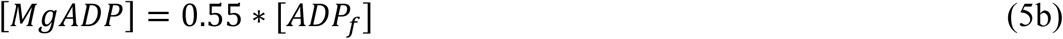

### Modelling changes in GK activity

The flux of glycolysis, which is limited by the rate of GK activity in the β-cell, is described as:

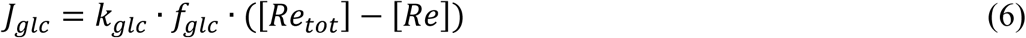

Where *k*_*glc*_ is the maximum rate of glycolysis (equivalent to GK activity), which was simulated as a normal distribution with a mean of 0.000126 ms^−1^ and standard deviation of 10% of the mean. [*Re*_*tot*_] = 10mM, the total amount of pyrimidine nucleotides. The ATP and glucose dependence of glycolysis (GK activity) is:

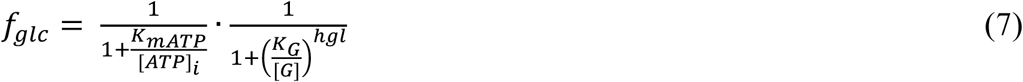

Where [*G*] is the extracellular concentration of glucose, *hgl* is the hill coefficient, *K*_*G*_ is the half maximal concentration of glucose, and *K*_*mATP*_ is the half maximal concentration of ATP.

GK deletion simulations, where GK was deleted in a population of cells, were modeled with a rate of glycolysis *k*_*glc*_= 0 ms^−1^ in randomly distributed cells. The number of cells that received the deletion was defined as the fraction *P*_*Mut*_ multiped by the number of cells (1000).

For GK inhibition simulations, decreases in *k*_*glc*_were modeled as

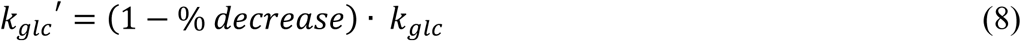

GK mutations that cause MODY-2 or PNDM were previously described in terms of their enzyme kinetics, 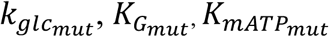 by [36,45-58], and were first simulated as:

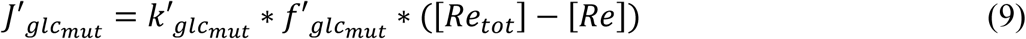

To account for variability in the control wild-type GK kinetics between studies (S8 Table), GK parameters 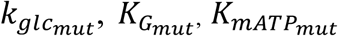 for each mutation were expressed relative to the wild-type GK parameter from the same study 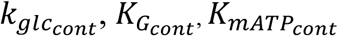. Thus:

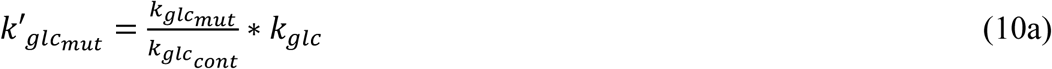

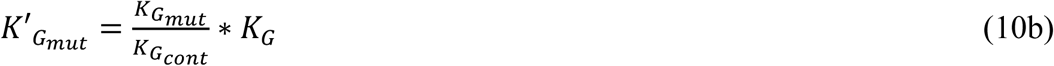

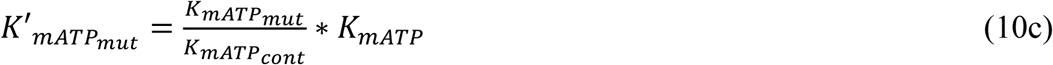

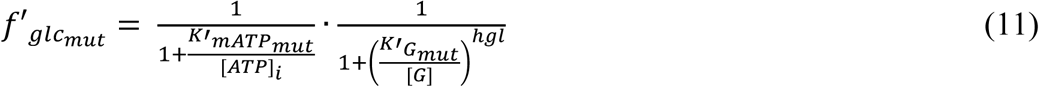

We also simulated MODY-2 mutants as a heterogeneous mutant:

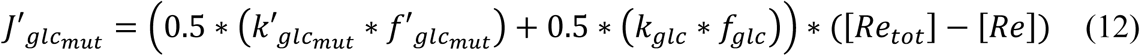

Where 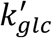 and 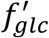 are described in (10a-10c, 11).

### Simulation data analysis

All simulation data analysis was performed using custom MATLAB scripts. The first 2000 time points were excluded to allow the model to reach a stable state. Cells were considered active if membrane potential, V_m_ exceeded −45mV. The fraction of cells that were active was determined relative to the total number of simulated cells (1000).

Duty Cycle was determined as the fraction of the [Ca^2+^] oscillations spent above a threshold value during the time course analyzed. This threshold value was determined as 50% of the average amplitude at 11mM glucose with no changes in rate of glycolysis. Duty cycle was reported as the average across all cells in the simulated islet.

Average parameters within a population of cells were calculated using a single simulation when the islet lacked electrical coupling (g_coup_=0pS) and when a fraction of the cells were active. This corresponds to either *P*_*Mut*_=0.50 (Fig.1) or *k*_*gle*_′ = (1 − .65) · *k*_*gle*_ (Fig.3). Each parameter was averaged over time where applicable, and across all GK^−^ cells/GK^+^ cells (Fig.1) or active/non-active cells (Fig.3).

When simulation results were binned according to OGTT values, ‘mild’ mutations had a 2h OGTT result of <145mg/dL, ‘moderate’ mutations had an OGTT result of 145mg/dL-175mg/dL, and ‘severe’ mutations had an OGTT value of >175mg/dL. Simulation results were also binned according to HbA1c and were characterized as mild if HbA1c <6.5%, moderate if HbA1c was 6.5-7.0% and severe if HbA1c >7.0%.

### Statistical analysis

All statistical analysis was performed in Prism (Graphpad). Either a Student’s t-test or a one-way ANOVA with Dunnett post-hoc analysis was utilized to test for significant differences between [Ca^2+^] activity, ΔNAD(P)H, plasma insulin, blood glucose, and simulation results. Matched pairs of Cx36^+/+^ and Cx36^−/−^ islets were chosen as islets imaged on the same day and a paired t-test was used to determine significance. If uneven numbers of islets with the same genotype were imaged on the same day, their results were averaged before the paired t-test was performed. Linear regression was used to evaluate the correlation between [Ca^2+^] activity, plasma insulin, or blood glucose vs. ΔNAD(P)H. Outliers were removed from experimental data sets based on the interquartile range (IQR). Data was first grouped by genotype, then outliers identified as any data point outside of [Q1 – 1.5*IQR, Q3 + 1.5*IQR] where Q1 and Q3 are the first and third quartiles, respectively. Data is reported as mean ± s.e.m. (standard error in the mean) or mean ± SD (standard deviation) where indicated. For data in Figure 5e and S6 some groups failed normality by an Anderson-Darling normality test (MATLAB) and therefore a non-parametric ANOVA (Kruskal-Wallis) and Dunn’s post-hoc analysis was used. Data is reported as mean ± s.e.m. unless otherwise indicated.

**Figure 1:**
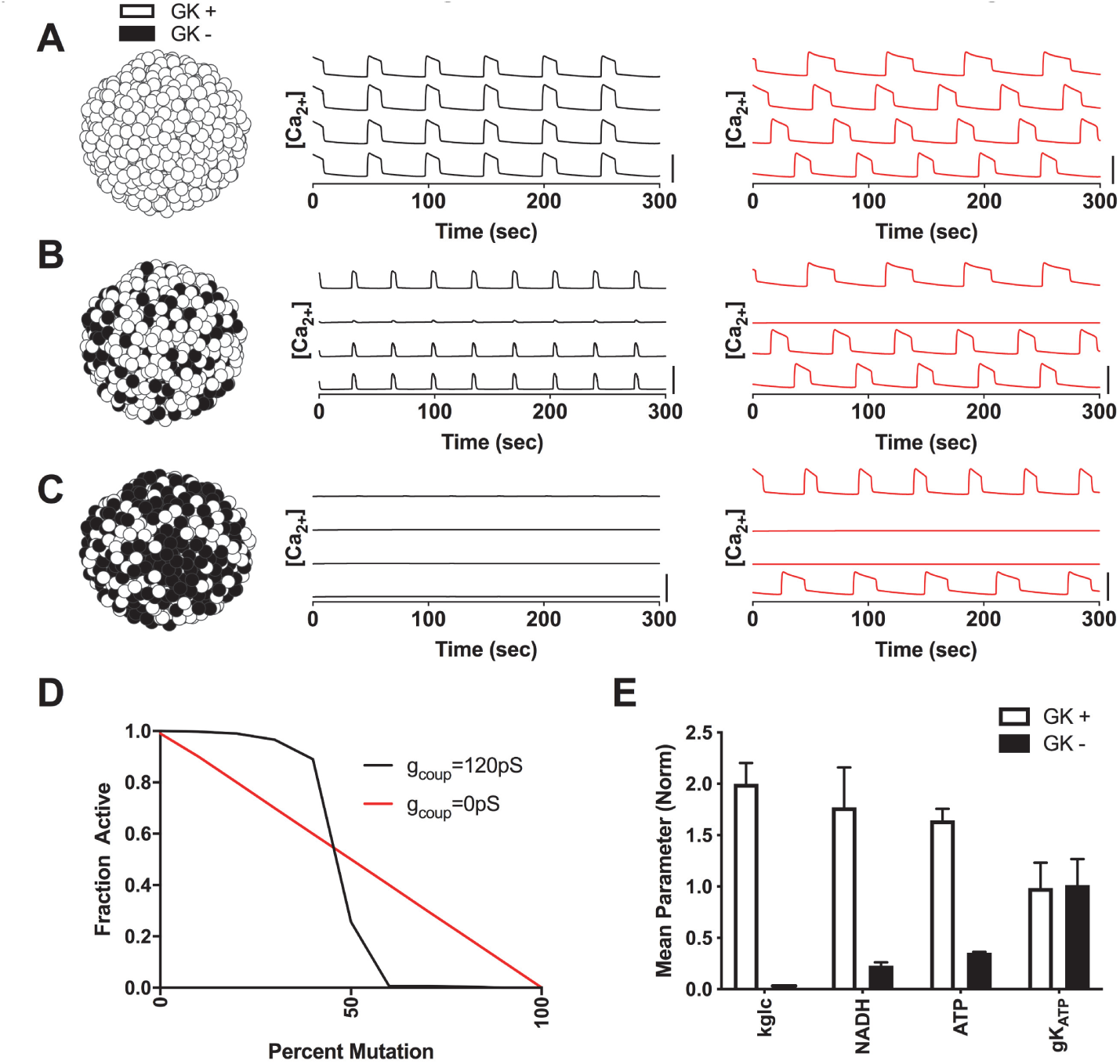
Simulating how metabolically deficient cells impact islet function via electrical coupling. A). Simulations of islet with normal GK activity. Left: Schematic of GK activity across simulated islet. Middle: Time courses of [Ca^2+^] for 4 individual cells within the simulated islet for g_coup_ = 120pS (black line). Right: Time courses of [Ca^2+^] for 4 individual cells within the simulated islet for g_coup_ = 0pS (red line). B). As in A. but for simulations where 40% of cells have no GK activity (GK-, black cells). C). As in A but for simulations where 60% of cells have no GK activity. D). Fraction of cells showing depolarization and elevated [Ca^2+^] activity (‘active cells’) in simulated islets vs the percentage of cells deficient in GK for g_coup_ = 120pS and 0pS. E). Average parameter values in simulation for cells with normal GK activity (GK^+^) compared to cells deficient in GK activity (GK^−^) at 50% GK^−^ cells. Error bars in E represent standard deviation over 500 cells. All simulations were run at 11mM glucose. Scale bar in A-C represents 0.5μM.

**Figure 2:**
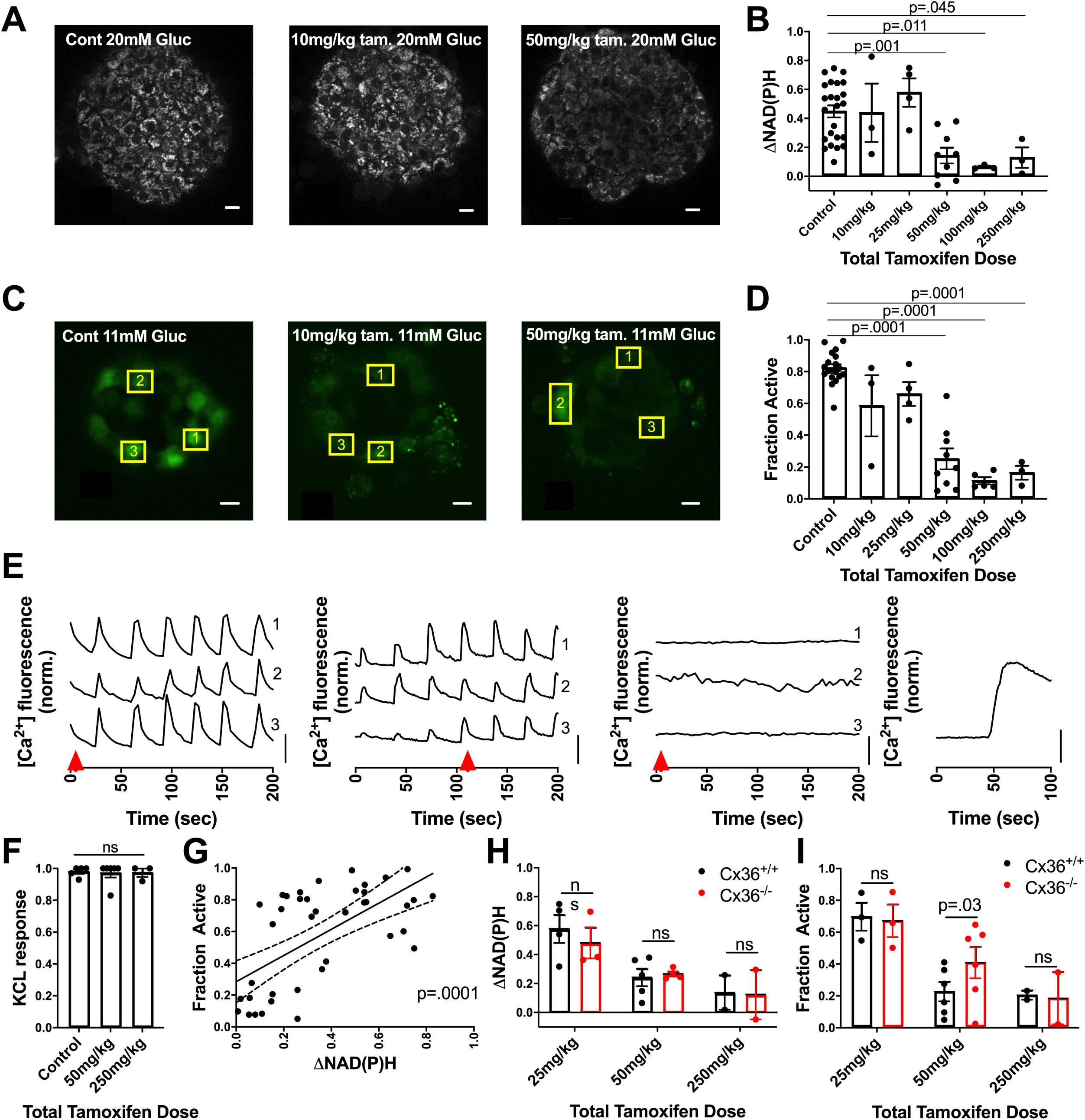
Experimentally demonstrating how metabolically deficient cells impact islet function via gap junction coupling. A). Representative images of NAD(P)H at 20mM glucose in islets from GK^lox/lox^;Pdx-Cre^ER^ mice injected with differing levels of tamoxifen, as indicated. B). Change in NAD(P)H from 2mM to 20mM (ΔNAD(P)H) compared to total tamoxifen dose. For control n=24 mice, for 10mg/kg n=3, for 25mg/kg n=4, for 50mg/kg n=9, for 100mg/kg n=3, for 250mg/kg n=3 (2-7 islets per mouse). C). Representative images of fluorescence from the [Ca^2+^] sensitive dye fluo-4 at 11mM glucose when injected with varying amount of tamoxifen. D). Fraction of cells showing elevated [Ca^2+^] activity (‘active cells’) at 11mM glucose compared to total tamoxifen dose. For control n=19 mice, for 10mg/kg n=3, for 25mg/kg n=4, for 50mg/kg n=9, for 100mg/kg n=5, for 250mg/kg n=3 (2-7 islets per mouse). E). Representative time courses of individual cells, shown by yellow boxes in C, at 11mM glucose within islets from mice treated with varying amounts of tamoxifen (From left to right: Control, 10mg/kg, 50mg/kg, 250mg/kg with KCl). F) Fraction of cells showing elevated [Ca^2+^] following KCl compared to total tamoxifen dose. For control n=5 mice, for 50mg/kg n=6, for 250mg/kg n=3 (2-4 islets per mouse). G). Scatter plot (dots) plus linear regression ±95% CI (solid and dashed lines) of mean active cells for islets from GK^lox/lox^;Pdx-Cre^ER^ mice vs mean ΔNAD(P)H for islets from the same mice (n=41 mice). p represents significance of linear trend slope. H). Comparison of ΔNAD(P)H for islets from GK^lox/lox^;Pdx-Cre^ER^;Cx36^+/+^ and GK^lox/lox^;Pdx-Cre^ER^;Cx36^−/−^ mice for the tamoxifen doses indicated. For 25mg/kg, n=4, for 50mg/kg n=5, for 250mg/kg n=2 (2-7 islets per mouse). I). Comparison of fraction of active cells at 11mM glucose for islets from GK^lox/lox^;Pdx-Cre^ER^;Cx36^+/+^ and GK^lox/lox^;Pdx-Cre^ER^;Cx36^−/−^ mice for the tamoxifen doses indicated. For 25mg/kg, n=3, for 50mg/kg n=6, for 250mg/kg n=2 (2-7 islets per mouse). Scale bar in A,C represents 10μm, scale bar in E represents dF/F of 50%.

**Figure 3:**
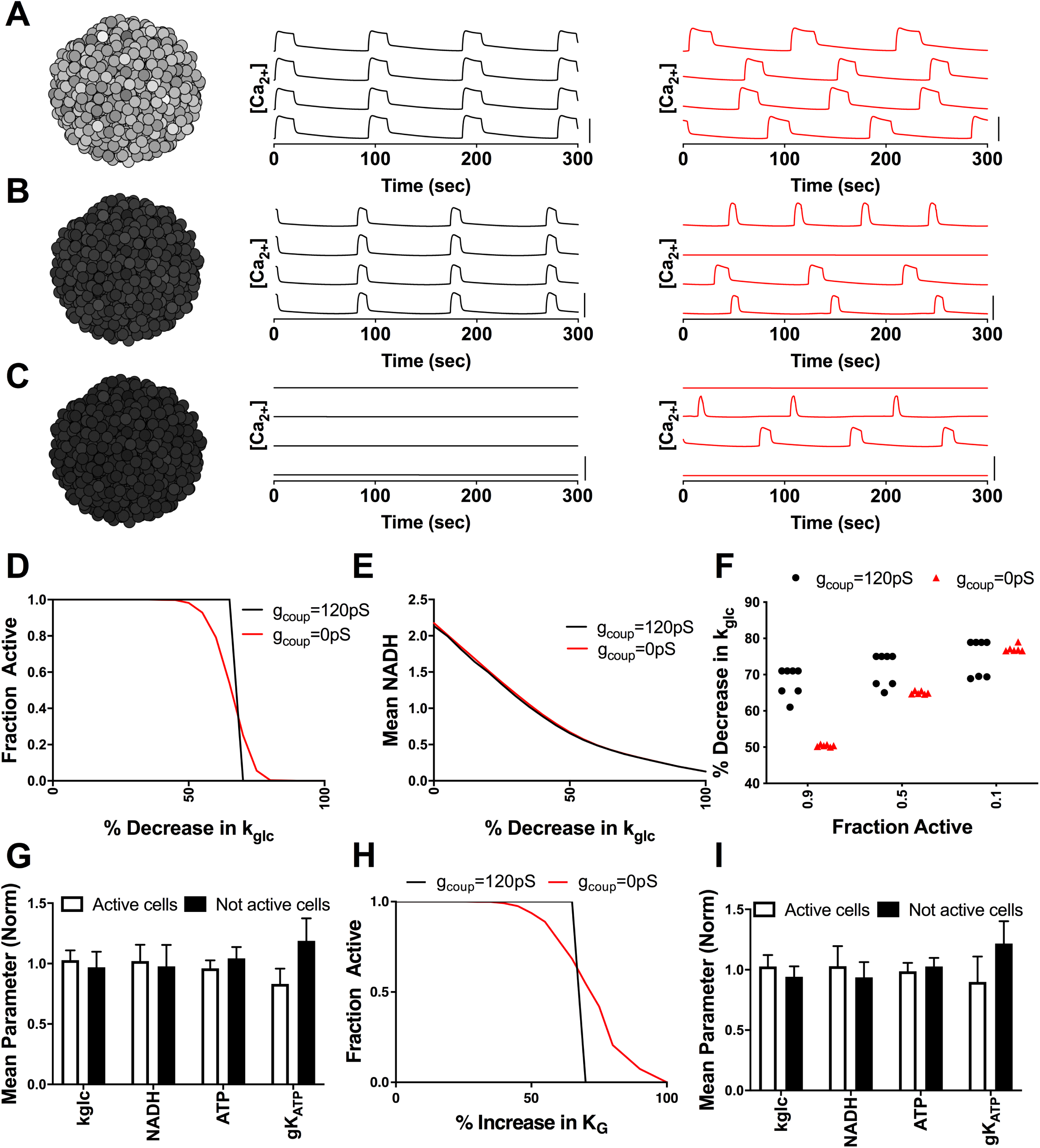
Simulations predicting how heterogeneity in metabolic activity impact islet function via electrical coupling. A): Simulation of islet with normal GK (k_glc_) activity. Left: Schematic of heterogeneous GK activity across simulated islet. Darker cells represent reduced GK activity. Middle: Time courses of [Ca^2+^] for 4 representative cells within the simulated islet for g_coup_=120pS. Right: Time courses of [Ca^2+^] for 4 representative cells within the simulated islet for g_coup_=0pS. B). As in A. but for a simulated islet with a 60% decrease in k_glc_ activity uniformly across the islet. C). As in A. but for a simulated islet with a 70% decrease in k_glc_ activity uniformly across the islet. D). Fraction of cells showing elevated [Ca^2+^] activity (‘active cells’) in a representative simulated islet vs. fractional decrease in k_glc_ (decreasing GK activity) for g_coup_ = 120pS and 0pS. E). Mean NADH across a representative simulated islet vs. fractional decrease in k_glc_ (decreasing GK activity) for g_coup_ = 120pS and 0pS. F). The decrease in k_glc_ required to reduce islet [Ca^2+^] activity to 0.9, 0.5, or 0.1 islets simulated with either g_coup_=120pS or g_coup_=0pS. Simulated data representative of 6 different random number seeds (g_coup_=0pS) and 7 random number seeds (g_coup_=120pS). G). Average parameter values in [Ca^2+^] active cells and non-active cells within a simulated islet with g_coup_=0pS and 65% decrease in k_glc_. H). Fraction of active cells, as in D vs. increases in the half-maximal concentration of glucose (K_G_), with g_coup_=120pS (black) and g_coup_=0pS (red).I) Average parameter values in [Ca^2+^] active cells and non-active cells as in G. with a 70% increase in K_G_. Data in G,I. represents mean ± SD across all active or inactive cells. Scale bar in A-C represents 0.5μM.

**Figure 4:**
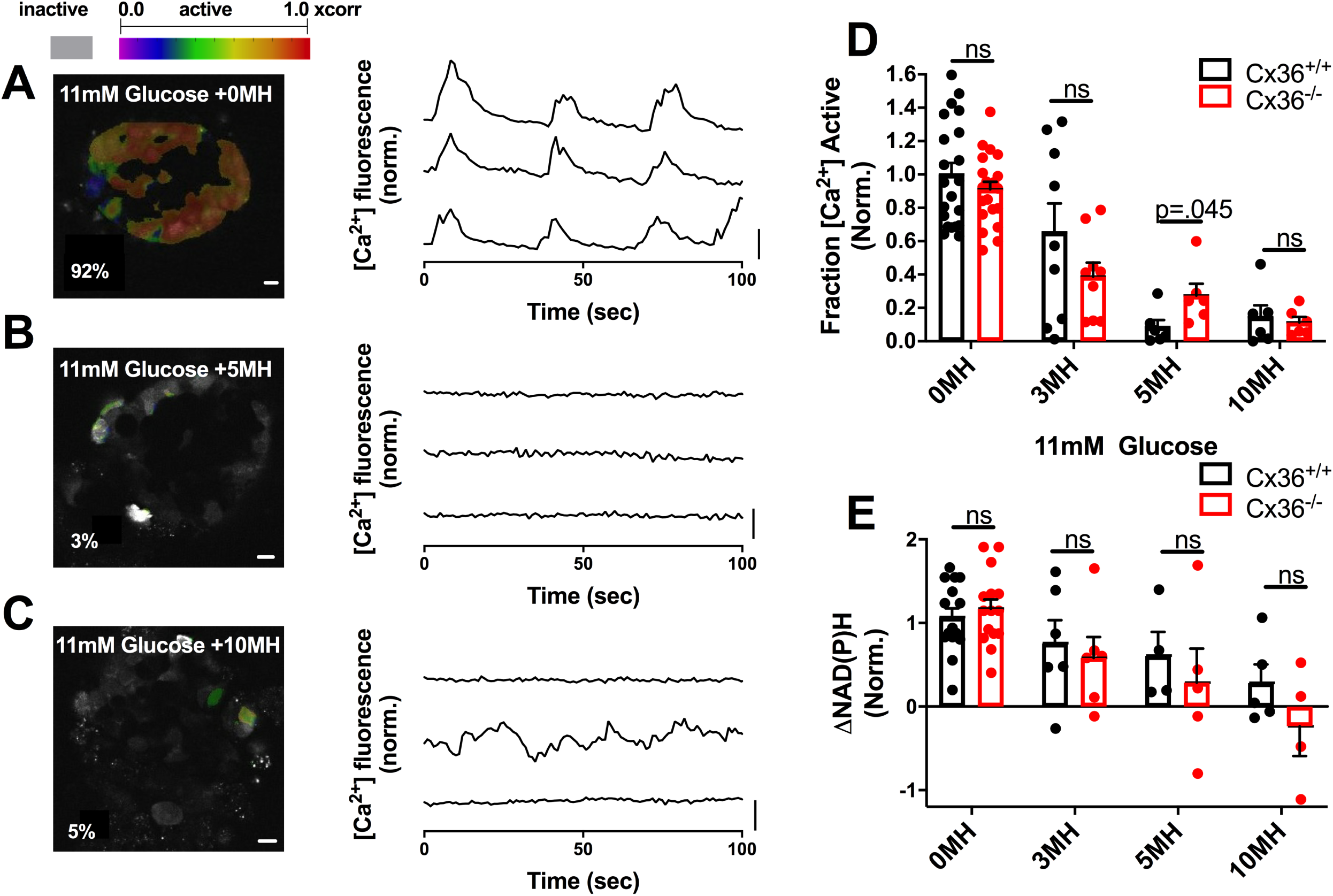
Experimentally demonstrating how heterogeneity in metabolic activity impacts islet function via gap junction coupling. A). Imaging [Ca^2+^] in islet at 11mM glucose with 0mM mannoheptulose. Left. False color map of regions of elevated [Ca^2+^] activity in islet Right: Time courses of [Ca^2+^] activity in 3 individual cells, as measured by fluo-4 fluorescence. B). As in A. but at 11mM glucose with 5mM mannoheptulose. C). As in A. but at 11mM glucose with 10mM mannoheptulose. D). Fraction of islet with elevated [Ca^2+^] activity with increasing mannoheptulose concentration for Cx36^+/+^ and Cx36^−/−^ islets. E). Change in NAD(P)H from 2mM to 20mM glucose with increasing concentration of mannoheptulose for Cx36^+/+^ and Cx36^−/−^ islets. For [Ca^2+^] activity experiments n=21 mice for 0mM MH, n=9 for 3mM, n=6 for 5mM MH, n=6 for 10mM MH (1-7 islets per mice). For NAD(P)H experiments n=15 mice for 0MH, n=6 for 3mM, n=5 for 5mM MH, n=5 for 10mM MH (1-6 islets per mouse). Data in D,E normalized to the mean [Ca^2+^] activity for 0mM MH. Scale bar in A-C images represents 10μm, scale bar in A-C time courses represents dF/F of 20%.

**Figure 5:**
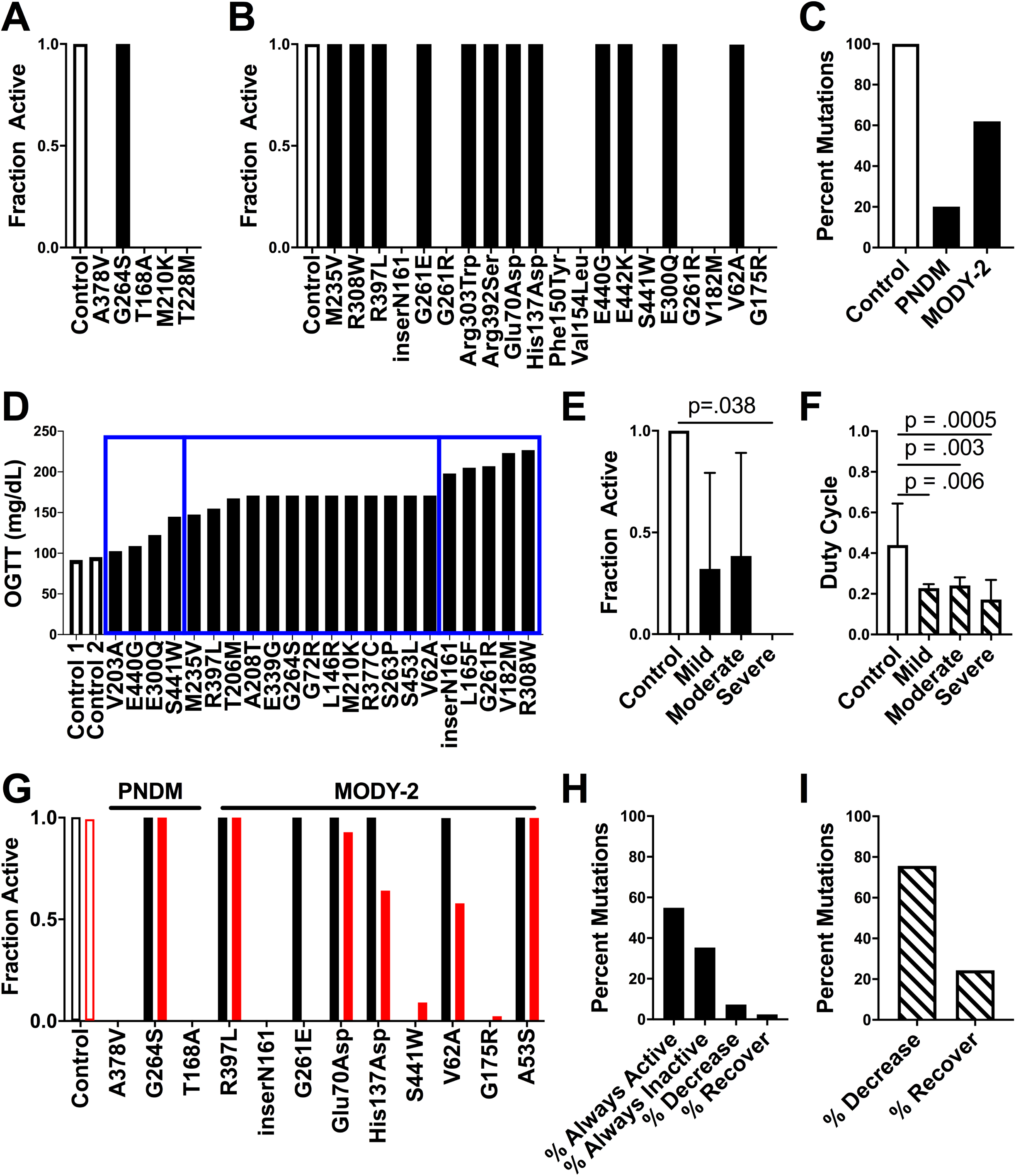
Simulations predicting how GCK mutations underlying monogenic diabetes impact islet function via electrical coupling. A). Fraction of cells showing elevated [Ca^2+^] activity (‘active cells’) in simulated islets from all PNDM mutations at 20mM glucose. B). Fraction of active cells in simulated islets for a representative subset of MODY-2 mutations at 20mM glucose. For all mutations see Fig.S2. C). Percentage of all mutations tested that had high islet [Ca^2+^] activity (>90% active cells) for Control (n=1), PNDM (n=5) and MODY-2 (n=82) at 20mM glucose. D). Oral glucose tolerance test results for a set of individuals with MODY-2 mutations, with bins indicating the severity of dysglycemia for mild, moderate, and severe mutations (left to right, blue rectangles). E). Mean fraction of active cells from simulations of mutations in D. at 11mM glucose, averaging over the disease severity and considering homozygous mutations. F). Average duty cycle for mutations in D. at 11mM glucose averaging over the disease severity and considering heterozygous mutations. All simulations A-F run with g_coup_=120pS. Data in E,F represents mean±SD. G). Fraction of active cells in simulated islets for a representative subset of PNDM and MODY-2 mutations with g_coup_=120pS (black) and g_coup_=0pS (red) at 20mM glucose. H). Percent of all MODY-2 mutations that maintain the same activity (always active, always inactive), decrease or increase (recover) [Ca^2+^] activity when g_coup_ is reduced from 120pS to 0pS, considering homozygous mutations at 20mM glucose. I). Percent of all MODY-2 mutations in which the duty cycle decreases or increases (recovers) when g_coup_ is reduced from 120pS to 0pS, considering heterozygous mutations at 11mM glucose.

## RESULTS

### Computational model predicts how discrete populations of metabolically deficient cells impact islet function

Previous studies have demonstrated how excitable and inexcitable cells (defined by K_ATP_ channel activity) interact within the islet via electrical coupling. Given that glucose metabolism varies extensively between previously identified β-cell sub-populations, we first set out to examine how cells with high and low levels of glucose metabolism interact. We simulated islets at elevated glucose with varying numbers of cells lacking GK activity. Simulated islets with normal electrical coupling and normal GK activity showed elevated [Ca^2+^] with coordinated oscillatory dynamics closely matching previously published models [11,22,42,59] (Fig. 1a). As the number of cells lacking GK activity (GK^−^) increased in number up to ∼40% of GK^−^ cells, the simulated islet retained normal coordinated [Ca^2+^] elevations albeit with a reduced plateau fraction (Fig. 1b). However above ∼50% GK^−^ cells, the simulated islet lacked elevation in [Ca^2+^] (Fig. 1c), with all remaining GK^+^ cells remaining silent. In the absence of electrical coupling, as the number of GK^−^ cells was increased, the disruption to [Ca^2+^] was more gradual (Fig. 1a-c), with all remaining GK^+^ cells responding. Thus, with fewer than ∼40% GK^−^ cells, the absence of electrical coupling diminished [Ca^2+^] compared to in the presence of coupling (Fig. 1d). However, with greater than 50% GK^−^ cells, the absence of electrical coupling increased [Ca^2+^]. Simulated GK^−^ cells showed reduced ATP and NADH levels, but normal total K_ATP_ conductance (Fig. 1e).

Thus, simulations predict the islet requires ∼50% of the cells to have near-normal glucose metabolism to show glucose-stimulated [Ca^2+^]. Below this number, the islet will sharply transition to lack any glucose-stimulated [Ca^2+^]. However, in the absence of electrical coupling the change in [Ca^2+^] is gradual and only cells with near-normal glucose metabolism will show glucose-stimulated [Ca^2+^].

### Experimentally testing how discrete populations of metabolically deficient cells impact islet function

To experimentally test these computational model predictions, we used a mouse model where GK deficiency is induced in a population of β-cells. We focused on testing: 1) That the threshold number of GK deficient cells required to sharply suppress islet [Ca^2+^] is ∼50%; and 2) That above this threshold number of GK deficient cells, a loss of gap junction electrical coupling elevates [Ca^2+^].

GK^lox/lox^;Pdx-Cre^ER^ mice received doses of tamoxifen totaling between 10mg/kg body weight (bw) and 250mg/kg bw. For tamoxifen doses 50-250mg/kg bw, blood glucose levels elevated significantly compared to control mice (see methods), with no significant difference in blood glucose between groups within this range (Supplemental Information, Figure S1). Below this dosage (10-25mg/kg bw) mice remained euglycemic, similar to control mice. Plasma insulin levels were significantly lower than control mice for tamoxifen doses 50-250mg/kg bw (Figure S1).

We isolated islets from these GK^lox/lox^;Pdx-Cre^ER^ mice and control mice that received varying tamoxifen dosages. We performed two-photon imaging of NAD(P)H to assess changes in the islet metabolic response (Fig. 2a). The NAD(P)H response decreased with increasing tamoxifen dosage (Fig. 2a), where doses 50-250mg/kg bw were significantly lower than in controls (Fig. 2b). This showed good correlation with blood glucose and plasma insulin (Figure S1). We also performed [Ca^2+^] imaging to assess changes in the islet electrical response (Fig. 2c). Doses of 10-25mg/kg bodyweight did not significantly impact [Ca^2+^], with coordinated [Ca^2+^] oscillations remaining showing varying patterns of slow or fast oscillations. In contrast, doses of 50-250mg/kg bw led to significantly lower [Ca^2+^] than in controls (Fig. 2d,e). However, the remaining [Ca^2+^] elevations were coordinated, indicating electrical coupling was not substantially diminished (Fig. 2e). For doses 50-250mg/kg bw KCl treatment elevated [Ca^2+^] across the majority of the islet, equivalent to in control islets (Fig. 2f), indicating that the disruption to [Ca^2+^] is due to an absence of metabolic-regulation of K_ATP_ and not due to cell death or de-differentiation.

The greatest decline in glucose-stimulated [Ca^2+^], from ∼70% to ∼20% of the islet showing elevated [Ca^2+^], occurred between 25 and 50mg/kg bw tamoxifen dosage. Given the lack of an associated fluorescent reporter, we were unable to directly quantify the % of cells showing GK deletion. However, the sharp decline in [Ca^2+^] occurred around an NAD(P)H response of approximately 0.15-0.2 (Fig. 2f), which is approximately 30-50% of the mean NAD(P)H response in control islets (Fig. 2b). This suggests that a sudden decline in [Ca^2+^] occurs upon a 50-70% loss in metabolic activity, which is equivalent to a complete loss of metabolic activity in 50-70% of cells. This is consistent with the simulated islet results (Fig. 1), where greater than 50% GK^−^ metabolically deficient cells leads to a dramatic decrease in [Ca^2+^].

We next tested whether a decrease in Cx36 and electrical coupling can increase [Ca^2+^] for levels of GK deletion above the threshold at which [Ca^2+^] is suppressed. We examined islets from age-matched GK^lox/lox^;Pdx-CreER;Cx36^+/+^ and GK^lox/lox^;Pdx-CreER;Cx36^−/−^ mice following tamoxifen injection. 50mg/kg bw tamoxifen is the smallest dose that led to markedly decreased [Ca^2+^] in the presence of Cx36 (Fig. 2d). At this dose similar NAD(P)H responses were observed in the presence and absence of Cx36, consistent with similar GK deletion (Fig. 2g). Despite this there was a significant elevation in [Ca^2+^] activity in islets lacking Cx36, consistent with simulation predictions (Fig. 2h). Islets from mice treated with either lower tamoxifen doses where islet [Ca^2+^] is near normal (25mg/kg bw), or higher doses where islet [Ca^2+^] is suppressed (250mg/kg bw), showed similar [Ca^2+^] irrespective of Cx36. This is also consistent with model predictions.

Therefore, islets lacking GK in a sub-population of metabolically deficient β-cells, show behavior in good agreement with computational model predictions. There exists a threshold number of ∼50% β-cells in which GK deficiency causes a sharp suppression in [Ca^2+^]; and this suppression depends on gap junction electrical coupling.

### Computational model predicts how a distribution of metabolically active/inactive populations impact islet function

We have examined how the islet responds with 2 distinct cell populations, with either elevated metabolic activity (normal GK) or deficient metabolic activity (deficient GK). While some studies have suggested distinct sub-populations exist within the islet with differing metabolic activity [19,25], other studies have demonstrated a continuous distribution of metabolic activity is present [29,31]. Within a continuous distribution, a ‘metabolically deficient’ population of cells may fall below some threshold metabolic activity required for stimulating [Ca^2+^]. We therefore simulated islets with a continuous distribution of GK activity between cells. With a normal distribution of GK activity, elevated [Ca^2+^] with coordinated oscillatory dynamics were observed in the presence of electrical coupling, but these oscillatory dynamics lacked coordination in the absence of electrical coupling (Fig. 3a). As the mean GK activity was reduced in the presence of electrical coupling, coordinated [Ca^2+^] elevations were maintained until a certain decrease in GK activity, at which point there was a transition to a complete absence of [Ca^2+^] elevation (Fig. 3b-d). In the absence of electrical coupling the decline in [Ca^2+^] was more gradual as GK activity was reduced (Fig. 3b-d). The change in NAD(P)H, indicating metabolic activity, was similar across the islet in the presence and absence of electrical coupling (Fig. 3e). As such, when the decrease in GK activity was low (<70% decrease), and the islet was metabolically active, the absence of electrical coupling decreased [Ca^2+^] as some cells lacked elevated [Ca^2+^] (Fig. 3d,f). However, with greater decreases in GK activity (>70%) such that the islet was metabolically deficient, the absence of electrical coupling increased [Ca^2+^] as some cells maintained elevated [Ca^2+^]. In the absence of electrical coupling, the population of cells where [Ca^2+^] elevations remained had slightly higher metabolic activity (higher k_gly_, NADH levels), and lower K_ATP_ open channel conductance (g_KATP_) (Fig. 3g) and thus were more excitable. When excluding heterogeneity in g_KATP_, similar observations were made when GK activity was reduced (Figure S2): the population of cells where [Ca^2+^] elevations remained had higher metabolic activity (higher k_gly_, NADH levels). Thus elevated metabolic activity was sufficient for the cells to be more excitable.

GK kinetics can be impacted in additional ways, such as substrate affinity [60]. To understand more generally how islet function can be impacted by changes in GK activity, we simulated islets with other perturbations to GK kinetics. When the affinity for glucose is reduced (increasing K_G_), the islet showed uniform elevated [Ca^2+^] until the affinity was reduced ∼2-fold (K_G_ increased by ∼70%), at which point [Ca^2+^] declined sharply (Fig. 3h). In the absence of electrical coupling the decline in [Ca^2+^] was more gradual (Fig. 3h), where the population of cells where [Ca^2+^] elevations remained had higher metabolic activity (higher NADH), and lower K_ATP_ open channel conductance (g_KATP_) (Fig. 3i). Reducing the affinity for ATP (by increasing K_mATP_) or increasing the hill coefficient for glucose affinity had much weaker effects on islet activity (Figure S3). Inclusion of stochastic noise under conditions of decreased GK activity also elevated [Ca^2+^] (Figure S3), consistent with prior predictions [26].

Thus, upon a distribution of cellular metabolic activity, similar results were observed as with 2 distinct populations of cells. The islet transitions from islet-wide [Ca^2+^] elevations to a complete lack of [Ca^2+^] according to the mean GK level: a lower mean GK level results in more cells falling into being metabolically deficient. The simulations specifically predict the islet requires >30% of the cells to have near-normal glucose metabolism to show islet-wide glucose-stimulated [Ca^2+^]. This behavior again depends on electrical coupling, where in the absence of electrical coupling the decline is gradual and excitable cells with near-normal glucose metabolism remain active.

### Experimentally testing how endogenous metabolically active/inactive populations impact islet function

To experimentally test these model predictions, we reduced GK activity across the islet with the GK inhibitor mannoheptulose (MH), in the presence (Cx36^+/+^) and absence (Cx36^−/−^) of gap junction coupling. In both the presence and absence of gap junction coupling, increasing concentrations of MH decreased [Ca^2+^] (Fig. 4a-c). In islets from Cx36^+/+^ mice the decline in [Ca^2+^] was modest for low MH concentrations (3mM), but a substantial decline occured between 3 and 5mM MH, and no further decline between 5 and 10mM MH (Fig. 4d). However, in islets from Cx36^−/−^ mice the decline in [Ca^2+^] was more gradual up to the maximal 10mM MH. At 5mM MH, where [Ca^2+^] was almost completely suppressed in islets from Cx36^+/+^ mice, [Ca^2+^] was significantly higher in islets from Cx36^−/−^ mice, where ∼30% of cells continued to show elevated [Ca^2+^] (Fig. 4d). For all MH treatments, the NAD(P)H response was similar between islets from Cx36^+/+^ mice and Cx36^−/−^ mice (Fig. 4e), indicating the differences in [Ca^2+^] are not a result of differing metabolic activity, which is consistent with simulated data (Fig. 3e).

Thus, good agreement with simulations that demonstrates a sharp decrease in [Ca^2+^] in the presence of gap junction electrical coupling as a result of decreased GK activity; but a gradual decrease in [Ca^2+^] in the absence of gap junction coupling. There is also good agreement with simulations, where 30-40% of cells maintain elevated [Ca^2+^] in the absence of gap junction coupling for a level of GK activity sufficient to suppress [Ca^2+^] in the presence of gap junction coupling.

### Computational predictions for gap junction coupling and *GCK* mutations that cause diabetes

Our results so far indicate that gap junction electrical coupling substantially impacts islet function when GK activity is heterogeneous. This includes enabling a large minority of metabolically active cells to increase [Ca^2+^] across the islet, and exacerbating the decline in [Ca^2+^] when a majority of cells show deficient GK and metabolic activity. We next applied our computational model to examine the role of electrical coupling in the presence of specific *GCK* mutations that cause NDM or MODY. We simulated the islet in the presence of electrical coupling and included altered GK kinetics based upon the biochemical characterization of a set of *GCK* mutations that cause MODY or PNDM (Table S1) [36,45-58]. The majority of PNDM *GCK* mutations (4/5) fully suppressed [Ca^2+^] at elevated glucose (Fig. 5a). In comparison, only a sub-set of MODY *GCK* mutations (∼35%) suppressed [Ca^2+^] at elevated glucose (Fig. 5b, Figure S4), showing more moderate impact than PNDM mutations (Fig. 5c). When accounting for MODY mutations being single allele variants, and thus modeling half of GK with normal kinetics, MODY *GCK* mutations reduced the [Ca^2+^] oscillation plateau fraction compared to controls (Figure S5).

We next compared simulation predictions for the impact of *GCK* mutations with corresponding clinical assessments for the loss of glucose control. We specifically examined a set of mutations for which patients had received oral glucose tolerance tests (OGTTs) and segmented the mutations into 3 classes (‘mild’, ‘moderate’ and ‘severe’) according to the level of glucose intolerance (2h blood glucose) (Fig. 5d). ∼50% of both mild and moderate mutations suppressed [Ca^2+^]. However, all severe mutations suppressed [Ca^2+^] (Fig. 5e). When accounting for MODY mutations being single allele variants, the [Ca^2+^] oscillation plateau fraction was reduced to a greater degree by severe mutations compared to mild/moderate mutations (Fig. 5f). Similar results in terms of suppression of [Ca^2+^] or impact on [Ca^2+^] oscillation plateau fraction were observed considering HbA1c, albeit with reduced difference between mild, moderate and severe mutations (Figure S6). These data show a good correlation between the clinical severity of *GCK* mutations and the simulated impact on [Ca^2+^].

With this agreement we examined the impact of electrical coupling on [Ca^2+^] upon *GCK* mutations. We simulated all MODY and PNDM mutations in the absence of electrical coupling and compared [Ca^2+^] to simulations in the presence of electrical coupling (Figure S4). No PNDM mutation showed a change in [Ca^2+^], but a subset of MODY mutations (∼10%) showed changes in [Ca^2+^] in the absence of electrical coupling (Fig. 5g,h). This impact of removing electrical coupling included increases in [Ca^2+^] for a subset of mutations in which [Ca^2+^] was normally suppressed; a reduction in [Ca^2+^] for different mutations in which [Ca^2+^] remained active (Fig. 5h,i). Thus, gap junction mediated electrical coupling plays a partial role in mediating islet dysfunction caused by *GCK* mutations.

## DISCUSSION

β-cells in the islet respond collectively to glucose, despite the individual β-cells themselves being highly heterogeneous. Prior studies have demonstrated that small changes in the oscillatory properties or glucose-responsiveness of individual β-cells can lead to large changes in the collective behavior of the whole islet [22,25,32,33]. Heterogeneity in metabolic activity between β-cells is extensive [28,29,31]. In this study we controlled the heterogeneity in GK activity between β-cells to examine how electrical coupling between metabolically heterogeneous cells impacts islet function. Using computational models and experimental systems, we discovered that small changes in the number of cells with deficient GK and glucose metabolism led to large changes in islet [Ca^2+^]. Electrical coupling was important, mediating either the recruitment of metabolically deficient cells to show elevated [Ca^2+^], or meditating the suppression of metabolically active cells to supress [Ca^2+^]. Based on these results, computational models further predicted a role for changes in electrical coupling to impact islet dysfunction in monogenic diabetes that results from mutations to *GCK*.

### Cell heterogeneity and phase transitions within the islet network

Our previous work examined the impact of heterogeneity in K_ATP_ channel activity in the islet, using computational models and experimental systems [22,61]. β-cells were rendered inexcitable via expression of mutant K_ATP_ channels that do not close in response to elevated metabolism, thus hyperpolarizing the cell. As the number of these inexcitable β-cells was progressively increased, the islet initially showed near-normal responses. However, at some critical number of inexcitable cells there was a sharp transition between islet-wide [Ca^2+^] elevations and complete suppression of [Ca^2+^]. Here, our computational and experimental results demonstrate similar findings. As the number of metabolically deficient cells, caused by reduced or absent GK activity, is increased beyond a critical threshold (40-50%) there is a sharp transition between islet-wide [Ca^2+^] elevations and complete suppression of [Ca^2+^]. The remaining cells with normal GK are suppressed via electrical coupling by the metabolically deficient cells. This is supported by results in the absence of coupling where remaining metabolically active cells show elevated [Ca^2+^] (Fig. 1). Our experimental results are consistent with these findings: the lowest level of GK deletion (in ∼50% of cells) sufficient to suppress islet [Ca^2+^] shows elevated [Ca^2+^] in the absence of gap junction coupling (Fig. 2).

We observed similar results whether GK activity was partially reduced across all cells in the islet (Figs. 3,4). Computationally (Fig. 3), there was a level of GK reduction at which there existed a sharp transition between islet-wide [Ca^2+^] elevations and complete suppression of [Ca^2+^]. This transition occurred where 30-40% of cells have sufficient metabolic activity to show [Ca^2+^] elevations if electrically isolated. Experimentally (Fig. 4), the lowest concentration of mannoheptulose that was sufficient to fully suppress islet [Ca^2+^] (5mM), only suppressed [Ca^2+^] in a sub-set of cells in the absence of gap junction coupling, leaving ∼30% of cells remaining active. The correspondence between computational and experimental results (Figure S7) indicates the presence of broad metabolic heterogeneity within the islet. Thus, when GK activity is partially inhibited, less metabolically active cells will have insufficient glucose metabolism to elevate [Ca^2+^], whereas more metabolically active cells will have sufficient glucose metabolism to elevate [Ca^2+^]. If there are sufficient numbers of cells capable of elevating [Ca^2+^] (predicted to be 30-50%, Figure S7), they can resist being suppressed by less metabolically active cells, and the islet can elevate [Ca^2+^]. These findings are summarized in Figure 6.

**Figure 6:**
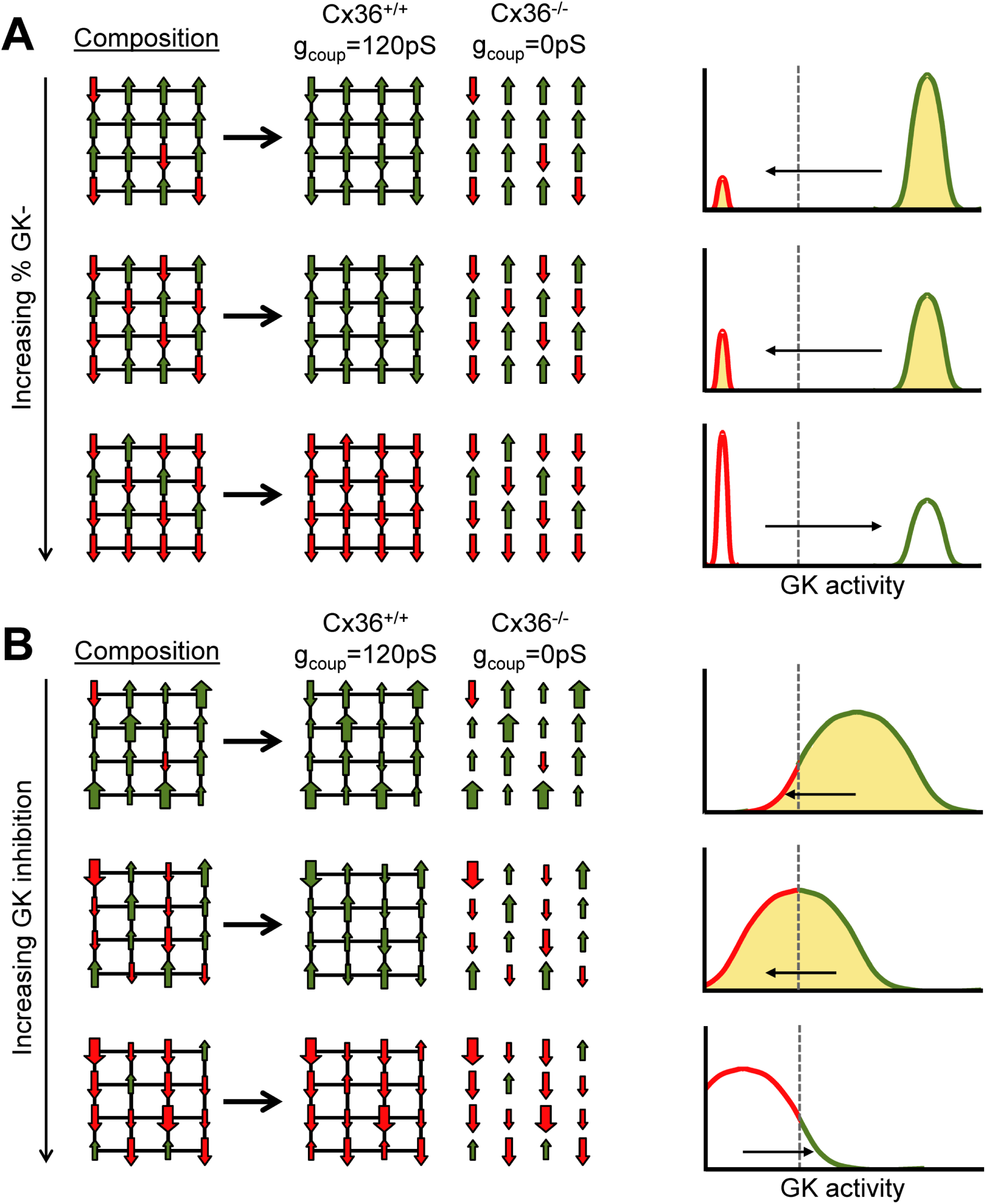
Model summarizing how cells higher in GK and lower in GK interact within the islets via gap junction electrical coupling. A) Schematic illustrating how composition of the islet under a bimodal distribution (as in Fig.1,2) impacts activity. Left: green-up represents metabolically normal cell that can elevate [Ca^2+^], red-down represents metabolically deficient cells that cannot elevate [Ca^2+^]. When deficient cells are in the majority electrical coupling cannot mediate the recruitment of elevated [Ca^2+^] (green). Right: equivalent histogram illustrating the same concept where shading represents recruitment and empty represents suppression. B) As in A for a continuous distribution of metabolic activity (as in Fig. 3-5). Large green-up represents highly metabolic active cells, large red-down represents highly metabolic deficient cell. Dashed line represents threshold GK activity for [Ca^2+^] elevation determining relative number of ‘metabolic deficient’ and ‘metabolic active’ cells.

Of note we do observe differences in this study compared to prior studies. Previously, when examining heterogeneity in K_ATP_ channel activity the number of inexcitable cells (cells with elevated K_ATP_ activity) required to suppress [Ca^2+^] across the islet was 20-40% (depending whether K_ATP_ overactivity was in a defined population of cells or broadly distributed [61]). This number is smaller than the number of metabolically deficient cells required to suppress islet-wide [Ca^2+^], which was 50-70%. Thus, populations of metabolically deficient cells are less effective at suppressing islet [Ca^2+^] than cells with elevated K_ATP_ activity. Conversely, this means smaller populations of metabolically active cells are effective at stimulating islet [Ca^2+^] which has significant implications for the role of metabolic heterogeneity in islet function (see discussion below). The residual ATP levels in metabolic deficient cells will partially close K_ATP_ channels, and therefore cells with elevated K_ATP_ activity may transmit greater hyperpolarizing current across the islet compared to metabolically deficient cells. Furthermore, in our model elevated ATP also regulates voltage-gated calcium channel (I_CaV_) activity, thus providing additional depolarization and Ca^2+^ entry at elevated glucose. Thus, cells with increased GK and metabolic activity may also transmit greater depolarizing current across the islet compared to cells with lower K_ATP_ activity. Since we were unable to precisely measure NADH changes on a cell-by-cell basis, we also cannot exclude the presence of metabolic coupling.

Therefore, following changes in glucose metabolism, a certain number of β-cells will show sufficient metabolism to elevate [Ca^2+^]. If this is above some critical number then the islet will elevate [Ca^2+^], whereas below this number a complete loss of glucose-stimulated [Ca^2+^] occurs. In the absence of electrical coupling, the population of β-cells with sufficient metabolism to elevate [Ca^2+^] will not be suppressed. These cells have sufficient GK activity for ATP generation and K_ATP_ closure, but may also have low K_ATP_ conductance, or other factors that promote elevated electrical activity and insulin release.

### Implications for Islet Physiology

It has long been known that β-cells in the islet are functionally heterogeneous [16], including heterogeneity in GK expression, glucose metabolism and mitochondrial function [28,30,31]. Recently using transcriptome, protein labelling, surface markers or fluorescent reporters, specific subpopulations have been defined with diminished GK expression or reduced mitochondrial function or glucose metabolism [27]. Our results indicate quantitatively how changes in the proportion of these cell sub-populations with deficient GK or glucose metabolism will impact islet function. Assuming the electrical coupling remains unchanged between sub-populations, the islet will remain tolerant to cells low in glucose metabolism and recruit these deficient cells to show elevated [Ca^2+^]. However, when they form a substantial majority (above 50-70%) islet electrical activity will be disrupted.

That the majority of β-cells need to be deficient in glucose metabolism for islet [Ca^2+^] to be suppressed means that gap junction electrical coupling generally serves to mediate the recruitment of metabolically deficient cells. Upon a minority of 30-50% of metabolically deficient βcells, gap junction electrical coupling mediates their activation by more metabolically active cells. Indeed, our computational and experimental results upon uniform changes in GK activity indicate that as few as ∼30% of metabolically active β-cells can recruit and elevate [Ca^2+^] across the islet (Figs. 3,4). Therefore, cells with high levels of glucose metabolism play an important role for elevating [Ca^2+^] across the islet via electrical coupling. As such the preferential disruption of metabolically active cell populations under pathogenic conditions could critically impact islet function.

Despite the import roll a large minority of metabolically active cells may play in elevating[Ca^2+^] in metabolically deficient cells, some studies have demonstrated that sub-populations low in glucose metabolism also show lower GJD2 expression (encoding Cx36) [19]. As a result, with increasing number of cells low in glucose metabolism, the islet electrical activity may trend towards the behavior we observe in the absence of electrical coupling. Cells low in glucose metabolism may not be recruited to elevate [Ca^2+^], and islet [Ca^2+^] elevations may remain in a population of metabolically active cells even if large numbers of cells are deficient in glucose metabolism. Thus, remodeling of electrical coupling may be protective against large numbers of cells with deficient glucose metabolism.

We simulated a Gaussian distribution of glucose metabolism which agreed well with our experimental results. However recent studies have indicated that small populations of cells, characterized by high electrical coupling and GK expression, play a substantial role in coordinating islet electrical activity [25]. Other highly metabolic subpopulations may also exist. Thus, within the populations of GK^+^ cells able to elevate [Ca^2+^] across the islet, a smaller sub-population may dominate this recruitment. Furthermore, cells with higher glucose metabolism may also show elevated gap junction coupling, again to aid in elevating [Ca^2+^]. Understanding the distribution of glucose metabolism, electrical coupling and other aspects of islet function will be important for future work.

### Implications for monogenic diabetes

Many forms of monogenic diabetes arise from mutations to genes that encode components regulating insulin secretion [62]. We previously applied computational models to demonstrate that eliminating electrical coupling could blunt the effect of many mutations to K_ATP_ channel subunits that cause NDM [26]. This relies on electrical coupling mediating the hyperpolarization and suppression of [Ca^2+^] by more inexcitable cells expressing mutant ATP-insensitive K_ATP_ channels. Our experimental and computational results show a role for electrical coupling in mediating islet-wide suppression of [Ca^2+^] by inexcitable cells that show decreased GK activity. For large decreases in GK activity (e.g. upon 5mM MH), a reduction in electrical coupling elevates [Ca^2+^]. Thus, we rationalized that for a subset of *GCK* mutations that cause diabetes, reducing electrical coupling may recover islet function.

Our computational model, which well described experimental data, also generated results that were consistent with clinical data regarding *GCK* mutations. Mutations that cause NDM suppressed [Ca^2+^] to a greater degree than mutations that cause MODY2, and those MODY2 mutations associated with poorer glucose control (OGTT or HbA1c) caused a greater reduction in [Ca^2+^]. Thus, we can be confident that the model can predict the effect of *GCK* mutations. The computational model predicts that reducing electrical coupling improves [Ca^2+^] elevations in a small number of mutations. This is consistent with the relatively narrow range of reduced GK or % cells low in GK over which electrical coupling mediates a suppression of islet [Ca^2+^] (Fig.1,3). The larger set of mutations in which electrical coupling promotes elevations in islet [Ca^2+^] is consistent with the idea that electrical coupling can mediate more metabolically active cells to elevate [Ca^2+^] in metabolically inactive cells (see discussion above). Thus, elevating electrical coupling may improve islet function in the face of *GCK* mutations, especially if electrical coupling is diminished by factors such as poor glucose control.

There are important considerations when modulating electrical coupling to promote β-cell [Ca^2+^] in the face of *GCK* mutations. Firstly, the terminal measurement we have made is glucose-regulated [Ca^2+^], which follows close agreement with our computational model. However a deficiency of GK and glucose metabolism will impact the production of mitochondrial derived amplifying factors that enhance insulin secretion [63]. Furthermore, paracrine signals that rely on metabolic products, such as glutamate, may also be disrupted and impact islet responses [64,65]. Therefore, while modulating gap junction electrical coupling can improve [Ca^2+^] upon GK deficiency, the elevation of insulin secretion may be modest given β-cells with low glucose metabolism would show low amplification. This may be overcome by stimulating incretin pathways such as GLP1R activation that amplify insulin secretion. We also note that our model is based upon a mouse islet and has been validated by mouse model data. The cyto-architecture and electrical regulation differs in human islets and there may be differences in GK action and electrical coupling in regulating islet function. However experimental data in human islets as to the role of heterogeneity and gap junction electrical coupling is limited and thus our ability to accurately simulate these properties and validate their role in the human context is limited. Nevertheless, our mouse-based computational model provided good agreement with clinical data, supporting our results and conclusions. Examining the role of heterogeneity in glucose metabolism and electrical coupling in regulating human islet function is an important goal for future work.

## CONCLUSIONS

The results from this study illustrate how gap junction mediated electrical coupling coordinate the response of metabolically heterogeneous β-cells within the islet. Using both experimental systems and computational models we have demonstrated that gap junction electrical coupling robustly coordinates the [Ca^2+^] response in the presence of high levels of metabolic heterogeneity. Electrical coupling promotes islet [Ca^2+^] elevations when low numbers of metabolically deficient β-cells are present but suppresses islet [Ca^2+^] elevations when higher numbers of metabolically deficient β-cells are present. Computational models support these findings, showing relevance to monogenic diabetes involving *GCK* mutations, where modulating electrical coupling can promote islet [Ca^2+^] elevations for certain mutations. Therefore, we provide further understanding how β-cell sub-populations regulate islet function via cell-cell communication.

## AUTHOR CONTRIBUTIONS

JMD designed and performed research, developed analytical tools, performed analysis, and wrote the manuscript. N.W.F.L. designed and performed research, developed analytical tools, and performed analysis. RAP performed research. WES performed research. OM performed research. MJW performed research and developed analytical tools. R.K.P.B verified results and wrote the manuscript.

## ACKNOWLEDGEMENTS

This work was supported by NIH grants R01 DK102950, R01 DK106412; and Juvenile Diabetes Research Foundation Grant 5-CDA-2014-198-A-N (to RKPB). Experiments were performed through the use of the University of Colorado Anschutz Medical Campus Advance Light Microscopy Core (P30 NS048154, UL1 TR001082); islet isolation was performed in the Barbara Davis Center Islet Core (P30 DK057516); and utilization of the JANUS supercomputer was supported by the NSF (CNS-08217944) and the University of Colorado. The authors would like to thank Dr Chris Rhodes for sharing the GK^lox/lox^ mice.

